# Parrots imitate song and dance with mixed vocal-gestural neural signals

**DOI:** 10.64898/2026.10.21.760654

**Authors:** Caleb Jones, Han Kheng Teoh, Julie Carpenter, Andrea Roeser, Ruidong Chen, Brian Kardon, Zhilei Zhao, Itai Cohen, Jesse Goldberg

**Author notes:** These authors contributed equally to this work (Caleb Jones, Han Kheng Teoh).

## Abstract

Humans shrug when saying “I don’t know” and nod when saying “yes,” yet it remains unclear how the brain coordinates vocal and gestural motor systems for learned, multimodal communication. Here we find that budgerigar parrots precisely imitate the fine temporal alignment between song and affiliative headbob gestures. Distinct colonies developed unique vocal-gestural motifs, which reorganized into new shared forms following social re-pairing. Neural recordings from the central nucleus of the anterior arcopallium (AAC), the cortical output of the parrot song system, revealed intermixed representations of vocalizations and head kinematics, including mixed selectivity at the level of single neurons. These findings show that socially learned multimodal communication is implemented by the joint representation of vocal and gestural motor signals within single neurons, revealing neural mechanisms that may extend to frontal cortical regions controlling speech and gesture in humans.

## Main Text

Multimodal coordination is a hallmark of complex social behavior. Humans spontaneously learn to couple speech with communicative gestures (*1*), yet the neural mechanisms that integrate distinct motor streams remain poorly understood. While songbirds have long served as a model for vocal learning, neural circuit-level studies have focused largely on the unimodal problem of matching vocal output to an auditory template (*2*, *3*). Consequently, it remains unclear how the brain coordinates vocal and non-vocal effectors for imitated multimodal communication.

While complex vocal-gestural coupling is widespread across animal species (*4–9*), domesticated budgerigar parrots (*Melopsittacus undulatus*) provide a unique opportunity to study displays mechanistically (*10–12*). In both natural and laboratory settings, budgerigars perform elaborate headbobs, rhythmic ∼5 Hz vertical movements during face-to-face warble singing (*10–12*). Originally rooted in motor patterns for juvenile allofeeding, budgerigars re-purpose headbobs in adulthood as an affiliative signal for social bond formation and maintenance (*11*).

Warble song is acquired through social imitation (*13*), yet it remains unknown whether the temporal coupling between vocalizations and communicative gestures is similarly learned. To test for social transmission of multimodal displays, we raised two colonies of budgerigars in acoustic and visual isolation from one another (Fig. 1A, Birds A1–A4 and B1–B3). Following a multi-month bonding period, we recorded synchronized vocalizations and body kinematics from individual birds using a directional microphone and high-speed camera positioned behind a one-way mirror (Methods). This experimental design permitted the isolation of single-subject behavior while maintaining social co-housing essential for species-typical behavior (Methods, 8.79 **±** 1.87 minutes and 1404 **±** 187 syllables per bird, n=7 birds).

**Fig. 1.**
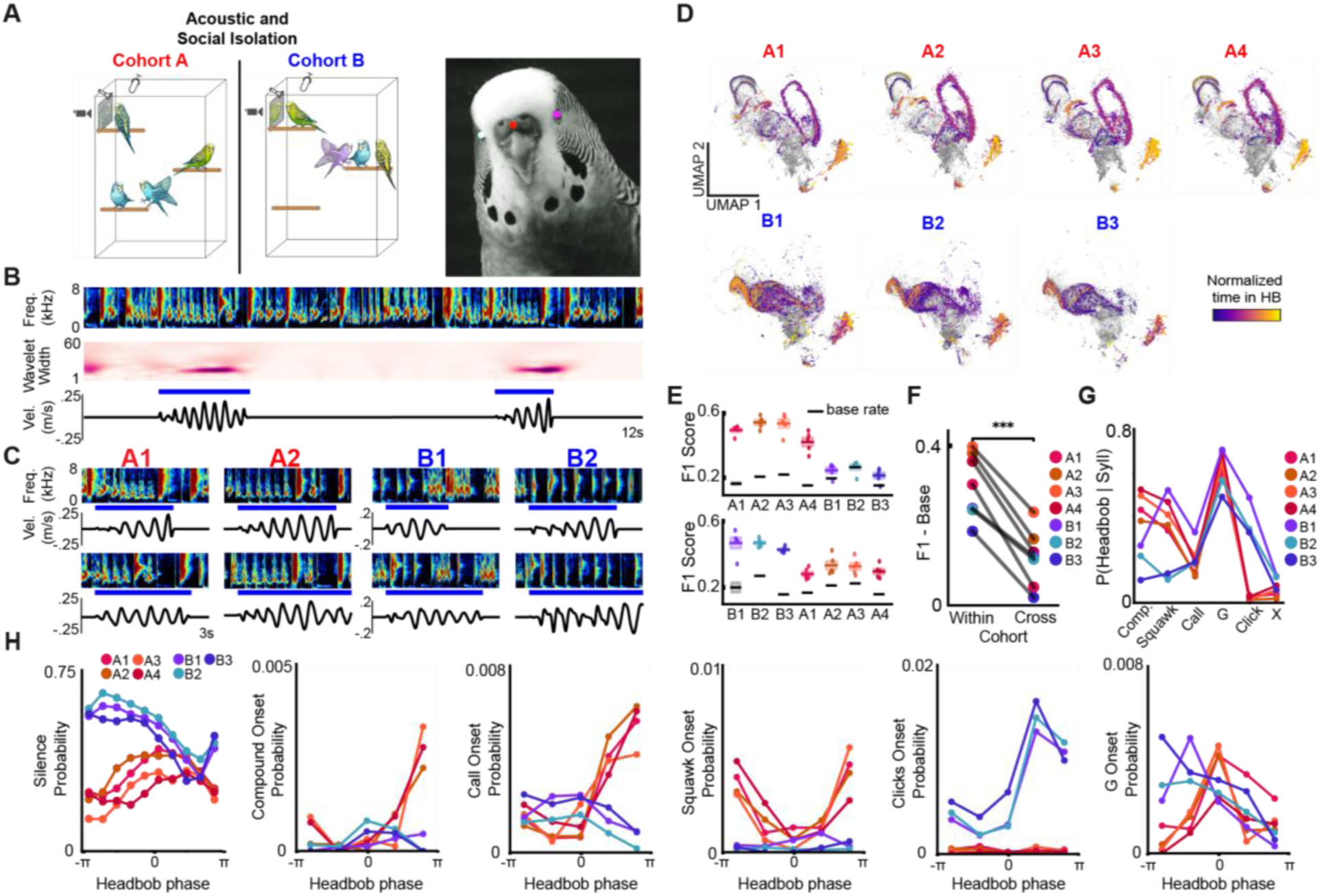
Distinct colonies exhibit distinct vocal-gestural coupling motifs. (**A**) Left: Schematic of homecages and cohort design; cameras and directional mics were positioned behind one-way mirrors at a remote corner of each cage. Right: representative video frame acquired from behind the mirror with DeepLabCut pose-tracking overlays. (**B**) Example song spectrogram (top), wavelet-based heat map of head displacement (middle; high-intensity regions indicate power used for automated headbob detection), and vertical head position trace with detected headbobs marked (red bars). (**C**) headbob onset–aligned spectrograms and vertical head position traces derived from DeepLabCut tracking. Cohort A birds preferentially couple compound syllables and squawks to headbobs, whereas Cohort B birds selectively couple clicks but not squawks. (**D**) UMAP projections of VAE acoustic latents for all birds (A1–A4, B1–B3), with points color-coded by sound timing relative to headbob onset (purple to yellow, onset to offset). Latent structure and temporal color clustering are shared within colonies but distinct across colonies. (**E**) Performance of LSTM_A1 (top) and LSTM_B1 (bottom) on predicting headbobs in heldout data from self and other birds. Each dot represents a distinct train/test split. For each bird the black bar indicates mean headbob rate, which sets chance LSTM performance, and each dot represents an independently train/test split for the LSTM. Note that LSTMs exhibit high within-cohort predictive power and low cross-cohort. (**F**) The average predictive performance of each bird’s LSTM was significantly higher within versus across cohort. (**G**) Probability of each syllable category being produced during headbobs. Colony B, but not A, bird click in their headbobs; all birds tend to end headbobs with the ‘g’ note, which contains a long low-pitch (<1kHz) tail (not shown) and occupies the lower right region of the UMAP projections in D. (**H**) Probability of silent gaps and initiation of distinct syllable types as a function of phase within the headbob cycle.

### Latent space analyses quantify warble song imitation

To quantify vocal imitation within social groups, we first used a supervised neural network (*14*, *15*) to segment songs into syllable categories established in past studies, including calls, clicks, squawks, compound syllables, and “G” notes (*16*, *17*) (fig. S1, see methods). Similarity analyses using conventional acoustic features such as pitch, entropy, and frequency modulation (*15*) confirmed imitation of syllable phonologies within colonies and divergence across colonies (fig. S1, see methods)(*13*, *17*).

Because syllable segmentation incompletely captures the fine acoustic structure of warble song (*18*), we next trained ‘shotgun’ variational autoencoders (VAEs) to analyze song as a continuous trajectory through acoustic latent space (*19*, *20*) (fig. S2, see methods). UMAP (*21*) visualizations of each bird’s acoustic manifold revealed striking within-colony imitation and cross-colony divergence (fig. S2D). To quantify these differences we derived a normalized similarity score from VAEs trained on individual birds, defined as the ratio of training-set to held-out error (see methods). This metric yields values near 1 for acoustically similar warble and progressively lower values for more divergent songs. A VAE trained exclusively on bird A1 warble (VAE_A1_) reconstructed data from birds A2-A4 with similarly low error as heldout data from A1, indicating shared warble acoustics in cohort A. Yet VAE_A1_ exhibited significantly higher error rates when reconstructing warble songs from birds B1-B3, revealing different acoustic structures. We repeated this pairwise analysis for VAEs trained on each bird individually and observed that networks reliably reconstructed cagemates’ warble with comparably low error but exhibited higher error for birds from the other colony, providing an independent quantification of within-colony convergence and cross-colony divergence of fine subsyllabic acoustic structure (Fig. S2B-C).

### Distinct colonies exhibit distinct vocal-gestural coupling motifs

We next wondered if vocal structure was temporally coupled to headbob kinematics (*10*, *11*). Using markerless pose tracking (*22*) and wavelet-based analyses, we identified headbob bout onsets and offsets across the dataset (Fig. 1B–C, see methods). Within headbob bouts, we defined each cycle as the interval between head position peaks, where −π and π represent consecutive peaks.

Multiple observations suggested that vocal-gestural coupling patterns were socially acquired. First, distinct syllable types were selectively coupled with headbobs in a cohort-specific manner. For example, in cohort A headbobs were associated with compound syllables but not clicks, a pattern reversed in cohort B (Mann–Whitney U test, p < 0.05; Fig. 1G). Second, the fine-scale timing of syllables was coordinated with the headbob phase in colony-specific ways. Colony B birds initiated cycles with silent gaps, producing a tendency for silence at the head’s most vertical position, whereas this tendency was absent in colony A (Fig. 1H; bird-level permutation test, p<0.05; Supplementary Videos 1-4; Methods). Third, cohort-specific preferences emerged for syllable category placement within the headbob cycle: colony A birds preferentially initiated calls and compound syllables near the end of headbob cycles and G syllables mid-cycle, whereas colony B birds preferentially initiated clicks near the end of the cycle (Fig. 1H). Notably, coupling did not appear to arise from fixed biomechanics. Head velocity did not consistently correlate with sound amplitude across birds and instead exhibited colony-specific patterns (fig. S3). Moreover, syllable types with common phonologies could be produced either inside or outside of headbob cycles (Fig. 1G), indicating that vocal content was flexible and not obligatorily constrained by head movement mechanics.

To visualize vocal-gestural coupling at subsyllabic resolution, we overlaid each individual bird’s acoustic manifolds with its headbob kinematic data. Each point in acoustic space (derived from the latent acoustic embedding from a single 80-ms spectrogram window, Methods) was labeled gray if produced outside of a headbob bout, or color-coded purple to yellow by latency from headbob onset if produced within a bout (Fig. 1D; Methods). If headbobs were randomly coupled to song, color-coded points would be uniformly distributed across the acoustic manifold. Instead, time-within-headbob bouts clustered in specific regions of acoustic space. Strikingly, birds A1–A4 and B1–B3 exhibited seemingly identical manifolds within cohorts, but distinct patterns across cohorts. These analyses show that coupling patterns were complex yet highly consistent within cohorts and divergent across cohorts.

To statistically test if finescale subsyllabic acoustic structure was coupled to headbob production, we trained Long Short-Term Memory (LSTM) networks to predict the timing of headbobs based on latent acoustic trajectories alone (Fig. 1E, Methods). Prediction performance was quantified using the F1 score, which ranges from 0 (no correct predictions) to 1 (perfect prediction with no false positives). Because headbobs were probabilistically coupled to syllable types, which could be produced both inside and outside of headbob bouts (Fig. 1G), F1 scores were expected to peak around ∼0.6, reflecting the empirically measured coupling probability in the labeled data (Fig. 1G). First, an LSTM trained to predict headbobs in bird A1 from a subset of its own warble song (LSTM_A1) reliably and accurately predicted headbob occurrence in held-out A1 data, indicating consistent within-bird vocal–gestural coupling. Strikingly, LSTM_A1 was equally effective at predicting headbob timing in birds A2–A4, despite never being trained on cagemate songs or head kinematics. Yet importantly, LSTM_A1 failed to predict headbob occurrence in birds from cohort B. Conversely, an LSTM trained on data from bird B1 (LSTM_B1) predicted headbob timing in cagemate B birds but significantly worse in A birds (Fig. 1E). Extending this analysis, we trained LSTMs on each individual bird and performed pairwise predictions across all bird pairs. This revealed a consistent pattern where vocal-gestural coupling generalized robustly within cohorts but not across cohorts (Fig. 1F). These analyses suggested that these fine-scale vocal-gestural motifs were learned during prior within-colony social bonding.

### Budgerigars socially learn vocal-gestural coupling patterns

To directly test if vocal-gestural coupling patterns are affected by new social interactions, we merged birds from cohorts A and B into dyads and allowed newly formed pairs to bond for at least 2 months (Methods). Both warble structure and its temporal locking to headbobs changed in each bird (Fig. 2). Plasticity in headbob motifs was visible in headbob-labeled acoustic manifolds (Fig. 2A–B, Supplementary videos 5-6). Consistent with these observations, an LSTM trained to predict bird B1’s headbobs from its warble after pairing with A1 (LSTM_B1_A1_) accurately predicted headbob timing in held-out data from B1_A1_. Remarkably, LSTM_B1_A1_ predicted coupling in the new cagemate A1_B1_ significantly better than in B1’s former cagemates (B2-B4) and even on B1’s own pre-pairing data (Fig. 2C). Extending this analysis, we trained LSTMs to perform all possible pairwise predictions across pre- and post-pairing datasets. Across all dyads, these analyses revealed that new vocal-gestural motifs emerged following pairings that were shared between current cagemates yet distinct from both former cagemates and even each bird’s past version of self (Fig. 2D). We noticed that B birds tended to evolve towards A bird patterns. To quantify this plasticity at the cohort level, we trained an LSTM classifier using vocal-gestural data from each post-pairing B bird alone and evaluated its out-of-sample generalization to pre-pairing cohort A and B birds. Prior to pairing, models generalized better within cohort B than to cohort A, reflecting a shared within-cohort pattern. After pairing with an A partner, this pattern reversed: the same models now generalized better to cohort A than to cohort B (Fig. 2E). This sign flip was consistent across independent training repetitions and observed for each B bird (B1–B3), indicating a cohort-level tendency of B birds to evolve towards A bird patterns (Fig. 2E). Together with the cohort-level differences prior to dyadic paring, these data show that budgerigars socially learn vocal-gestural coupling.

**Fig. 2.**
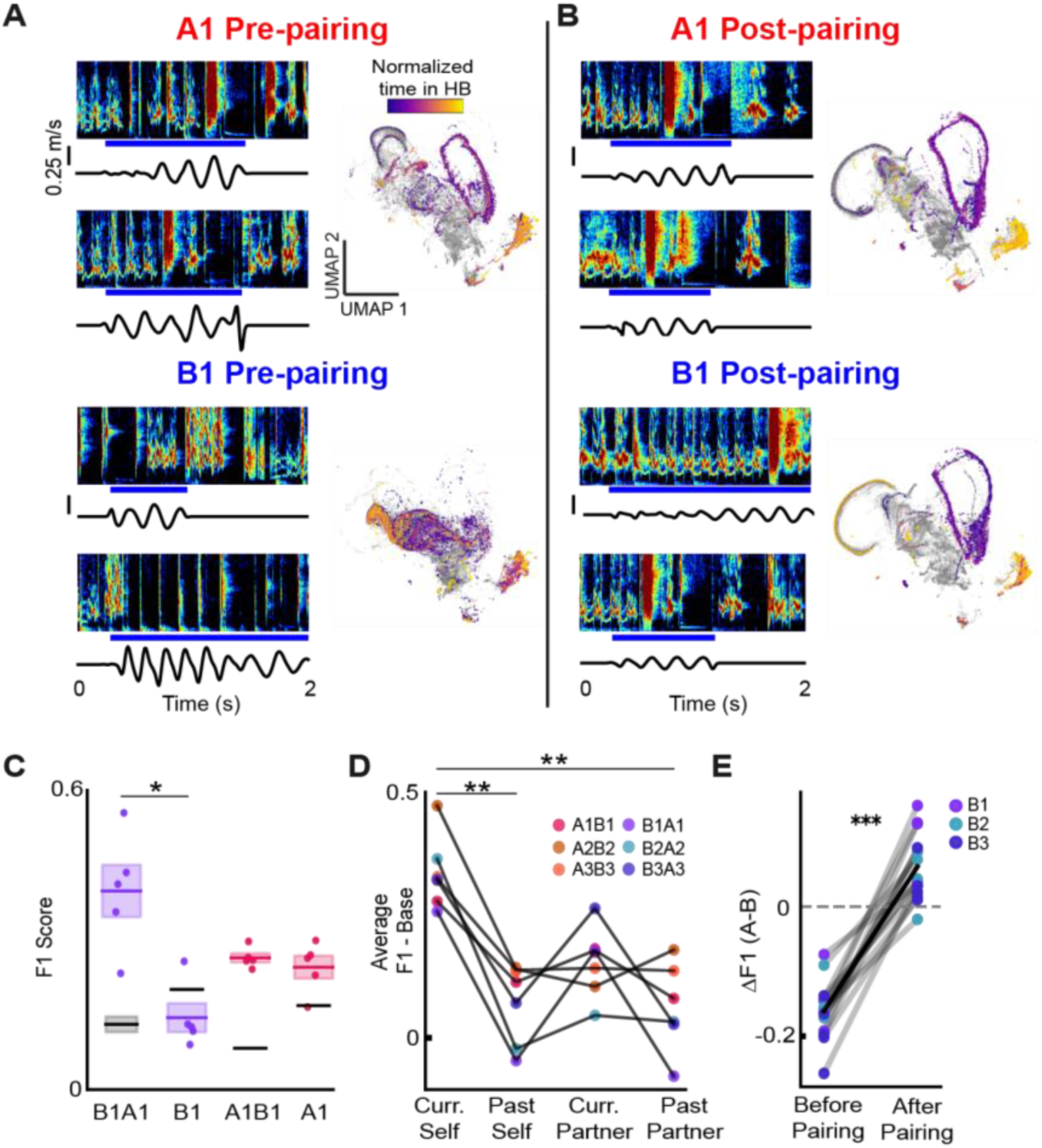
Budgerigars socially learn vocal-gestural coupling patterns. (**A**) headbob onset–aligned spectrograms and vertical head velocity traces for birds A1 and B1 prior to pairing. A1 preferentially produces chortles and squawks during headbobs, whereas B1 produces clicks. Right: UMAP projections of acoustic latent embeddings for A1 and B1, colored by normalized position within the headbob bout (purple, early; yellow, late). (**B**) Same analyses after 7 weeks of pairing. B1 adopts A1’s chortle–squawk pattern during headbobs, and post-pairing latent manifolds converge, consistent with socially driven plasticity. (**C**) LSTMs trained on post-pairing B1 vocalizations to predict headbob occurrence are evaluated on held-out B1 post-pairing data, B1 pre-pairing data, B1 post-pairing data, and A1 pre-pairing data. Models generalize better to the current cagemate than to the same bird prior to pairing. Black bars indicate baseline (chance) headbob rate; dots represent cross-validation folds. Asterisks represent significant results of a paired t-test, *p<0.5, **p<0.01 (**D**) Summary of LSTM performance across birds, shown as mean improvement in F1 score over baseline for models trained post-pairing and tested on current self, past self, current partner, and partner pre-pairing vocalizations. Bird identities are labeled by colony and pairing partner (e.g., A1B1 denotes bird A1 after pairing with B1; conversely B1A1 denotes bird B1 after pairing with A1). Asterisks represent significant results of paired t-test, **p<0.01 (**E**) Change in LSTM performance for predicting headbobs of new A cagemates using models trained on each B bird’s vocalizations before versus after pairing.

### Vocal and gestural representations are spatially intermingled within AAC

Precise coupling of vocalizations to headbob and headbob phase requires neural signals driving both vocalizations and headbob kinematics to be tightly coordinated. This could be implemented by intermixing of premotor signals in forebrain nuclei of the parrot song system. Alternatively, gesture and vocal premotor systems could remain segregated throughout the forebrain (*23*, *24*) and only coordinate at brainstem structures known to mediate unlearned respiratory-vocal-gesture coupling in bats, rodents, suboscines and fish (*25–27*). Like songbirds, parrots have a nucleated song system (*28*, *29*). AAC, the cortical output of the parrot song system, is necessary for song production and exhibits song-locked activity, including correlations with fine acoustic structure (*30*). Unlike songbird RA, which exhibits limited connectivity with surrounding tissue and which projects specifically to respiratory and vocal motor nuclei of the brainstem, AAC has dorsal ‘core’ and ventral ‘shell’ regions that are reciprocally interconnected with each other, with surrounding shell tissue, and with other thalamic and cortical nuclei of the song system (*28*, *29*). To sample neural representations in the entire AAC region, we chronically implanted moveable electrode arrays targeted dorsal to AAC and advanced electrodes ∼1.5 mm through dorsal core and ventral shell regions (fig. S4). We recorded vocalizations, neural activity and head movements with skull-mounted accelerometers during 1-2 months of natural social interactions (see methods). We advanced electrodes through both structures over 2 months of recordings and recorded 271 neurons in 4 birds. Headbobs produced during warble song produced pronounced oscillations in the accelerometer signal, enabling detection of HB onset and phase (Fig. 2, see methods).

We aligned AAC activity to WhisperSeg-labeled syllables and found that most neurons exhibited firing rate modulations associated with vocalizing. In total, 211/271 neurons showed a significant firing-rate maximum or minimum between 100 ms before syllable onset and the median syllable duration. These neurons exhibited diverse patterns of singing-related activity, including phasic activations or suppressions at syllable onset (figs. S4-5). In many cases, activity preceded syllable onset and correlated with upcoming acoustic features such as pitch and amplitude, consistent with a premotor role (fig. S5). These vocalization-related signals are consistent with a recent study recording from AAC, suggesting that we sampled a similar neuronal population (*30*). Vocalization-locked activity was distributed throughout dorsal and ventral AAC without spatial bias (Fig. S4; p=0.43, permutation test of depth effects using a variance-weighted PCA linear mixed-effects model, see methods).

The syllable-level coordination between vocalizations and headbobs, and their millisecond-scale coupling to headbob phase, requires premotor vocal signals to have information about headbob kinematics. To test if AAC activity was associated with headbobs, we aligned spiking to headbob onsets and to the phase of individual headbob cycles and found that a large fraction of neurons exhibited headbob-locked activity at recording sites that overlapped with vocal representations (Fig. 3; fig. S3). Many neurons could exhibit activations or suppressions during headbob bouts (activated: 28, suppressed 59, Fig. 3). To examine cycle-locked activity, we expressed spike times relative to headbob phase and found that over half of AAC neurons showed significant phase preference (82/151; Rayleigh’s test, p < 0.01), collectively tiling the headbob cycle (Fig. 3E-G). PCA decomposition of headbob-aligned firing rates revealed no effect of recording depth, indicating that the same AAC population that carries vocalization-related activity also represents headbob onset and phase (fig. S4).

**Fig. 3.**
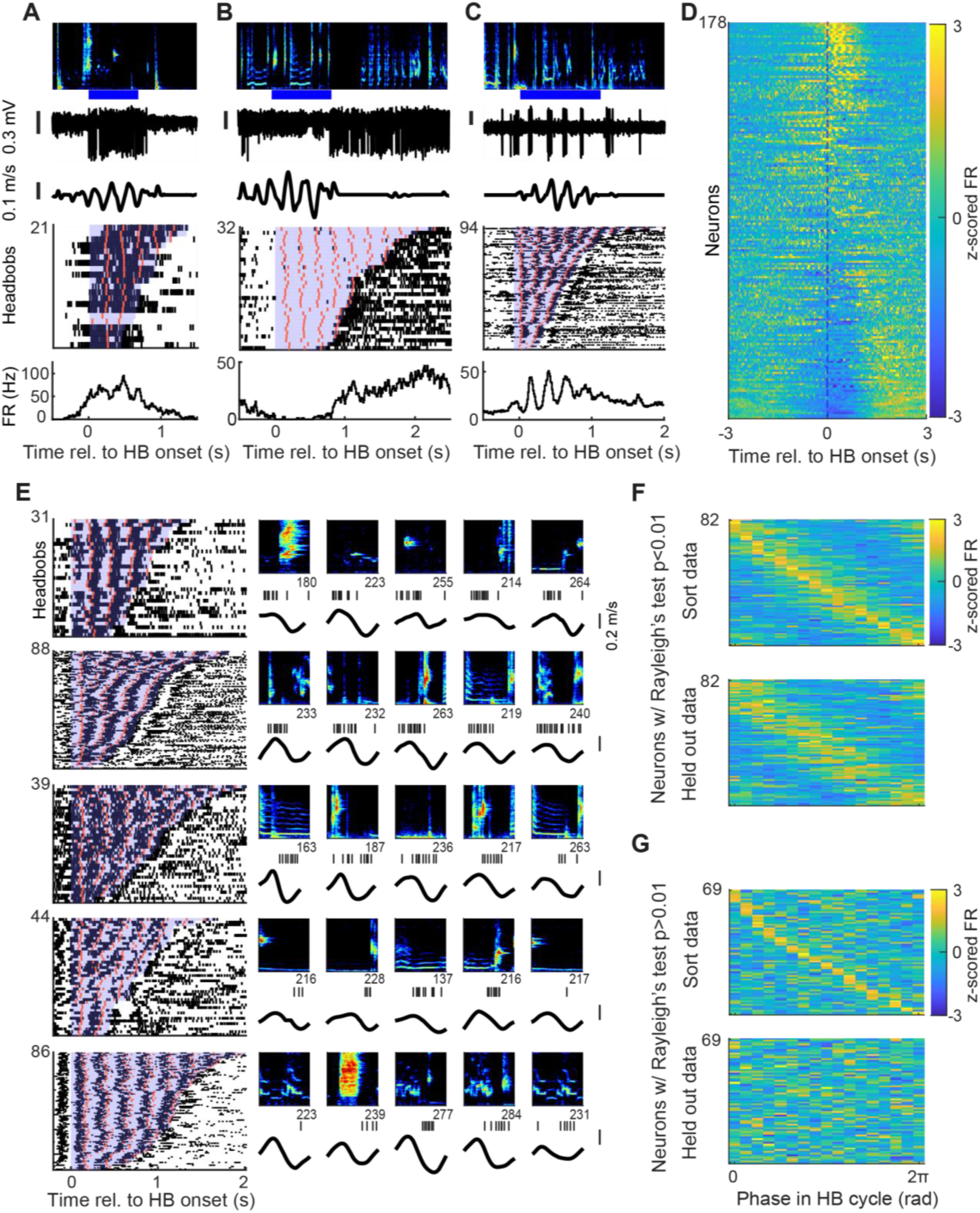
**AAC activity locked to headbob onset and phase**. (**A–C**) Top to bottom: headbob onset–aligned spectrograms, spike trains, vertical head velocity, rasters, and firing-rate histograms for three example neurons. Rasters sorted by headbob bout duration (blue shading); orange ticks indicate acceleration peaks within each headbob cycle. (**D**) headbob–aligned, z-scored firing rates of 178 neurons, sorted by the first principal component (PC1) score of population headbob–aligned activity. (**E**) Five example phasic neurons ordered by preferred headbob phase. Left: headbob–aligned raster plots as in A–C. Right: five example individual headbob cycles; from top to bottom, audio spectrogram, spike times (black ticks), and head velocity during each cycle. Scale bars, 0.2 m/s. Numbers below individual cycle spectrograms indicate duration in milliseconds. **(F**) As a population, AAC neurons tile headbob phase. Top: held-in headbob bouts used to determine sort order. Rows show neurons with significant Rayleigh test results, ordered by the phase of peak z-scored firing. Bottom: same neurons plotted for the held-out 50% of headbob bouts. The preservation of the cycle tiling in heldout data confirms that tiling is not the result of sorting artifact. (**G**) Same as F, but for neurons with non-significant Rayleigh test results, confirming that these neurons are not related to phase.

### Single AAC neurons exhibit mixed selectivity for vocalization and headbob kinematics

Because song and head kinematics were probabilistically correlated, apparent headbob-locked activity could arise even from purely vocal premotor signals. We used two complementary analyses to dissociate vocal and gestural contributions to AAC spiking. First, calls and squawks commonly occurred either during or outside headbob bouts in the recorded birds. Because these syllable types exhibit within-category acoustic stereotypy, this provided an opportunity to examine the influence of headbobbing on syllable-locked neural signals. For each call or squawk syllable rendition produced during a headbob, we identified its nearest acoustic neighbor produced outside of headbobs, providing trial-matched comparisons with highly similar acoustics. Despite acoustic matching, many neurons exhibited significantly different syllable-aligned firing when the same syllable was produced inside versus outside headbobs (fig. S6, 16/26 call-locked neurons and 13/30 for squawk-locked neurons, p<0.01, permutation test, see methods). Thus syllable-locked activity could be influenced by concurrent headbobs.

To more rigorously quantify the contributions of vocal acoustics and head kinematics to single-neuron activity, we used gradient-boosted decision tree ensemble models, XGBoost, to predict firing rates during headbob bouts (*31*, *32*). For each neuron with at least 20 headbob renditions, we predicted mean firing rate throughout the headbob in 50 ms bins using three models: vocal features only, movement features only, or a full model with both feature sets. Vocal features consisted of the low-dimensional latent representation of the acoustic signal, capturing fine-scale spectrotemporal structure, whereas movement features included instantaneous head velocities along orthogonal axes, headbob phase, and timing within a headbob bout (see methods). A null model predicted each neuron’s mean firing rate in each bin. Performance was quantified as improvement in mean squared error relative to the null model (%ΔMSE) and assessed using one-sided Wilcoxon signed-rank tests across held-out files.

Nested model comparisons were used to classify neurons. In full models, this approach powerfully predicted observed single neuron discharge (Fig. 4F, 59/138 neurons with significant full models, p<0.05, mean %ΔMSE=42.6, range: 11.3 to 84.2). Neurons were classified as purely vocal if removal of movement features did not degrade performance, but removal of vocal features did (F vs V, p>0.001 and F vs M, p<0.001, e.g. Fig. 4A, E, left, 30/59 vocal neurons). Conversely, neurons were classified as purely movement if removal of vocal features had no effect but removal of movement features significantly reduced performance (Fig. 4B, E, middle, p<0.001, 14/59 movement neurons). Neurons were classified as mixed if removal of either feature set significantly degraded model performance (Fig. 4C, E, right, 15/59 mixed neurons). Using this variance-partitioning framework, we identified neurons whose activity was explained predominantly by vocal acoustics, head kinematics, or jointly by both (Fig. 4G). Importantly, mixed neurons required the combination of acoustic and kinematic information to accurately predict spiking, demonstrating genuine mixed selectivity rather than indirect coupling via correlated features. Together these analyses show that AAC mixes vocal and gestural neural signals, even at the level of single neurons.

**Fig. 4.**
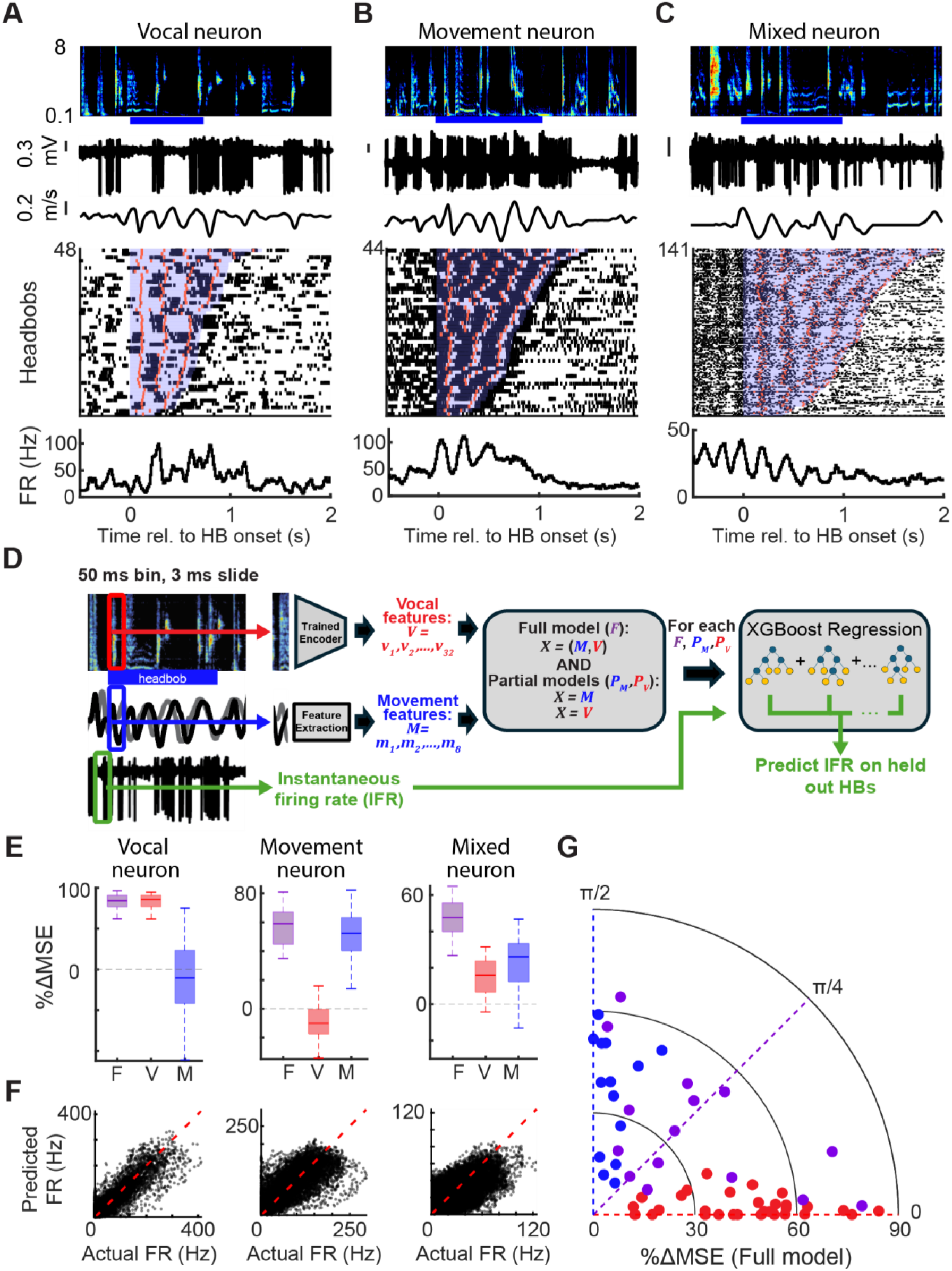
Single AAC neurons encode vocalizations, gestures, and both. (**A-C**) Example neurons classified by encoding models as vocal-only, movement-only, or mixed. From top to bottom: headbob onset–aligned spectrograms, spike trains, head velocity, spike rasters, and firing-rate histograms. Rasters are sorted by headbob bout duration (blue shading); orange ticks mark acceleration peaks within each headbob cycle. (**D**) Schematic of the gradient-boosted decision tree regression model (XGBoost) used to predict single-neuron firing rates from vocal and/or movement features. (**E**) Model performance for the neurons in (**A–C**), expressed as percent improvement over a null model. Performance is shown for the full (vocal+movement; F, purple), vocal-only (V, red), and movement-only (M, blue) models. Center lines indicate medians across 40 train/test splits; boxes show interquartile ranges and whiskers extend to 1.5× IQR. Dashed line indicates null-model performance. (**F**) Predicted versus observed firing rates for the full model for the example neurons in (A–C); dashed red line indicates unity. (**G**) Polar summary of encoding properties across neurons. Radius denotes full-model performance relative to null; angle denotes mixing index (0 = purely vocal, π/2 = purely movement). Each point represents one neuron (vocal, red; movement, blue; mixed, purple).

## Discussion

Human communication is multimodal but neural mechanisms for coordinating vocal and nonvocal effectors remain poorly understood. While vocal learning is typically studied as a unimodal auditory to vocal motor learning problem (*2*, *33*, *34*), our results demonstrate that budgerigars imitate both vocalizations and their precise coupling to communicative gestures. This “embodied singing” is highly plastic, with vocal-gestural motifs reorganizing to match new social partners.

At the neural level, neurons in AAC, the cortical output of the parrot song system, jointly represent syllable timing, fine-scale acoustics, and headbob kinematics. A substantial population of these neurons exhibited mixed selectivity: accurate prediction of spiking required the combined inclusion of both acoustic and kinematic variables. Mixed selectivity has been described in frontal cortical areas during cognitive tasks that require processing of multiple task variables for decision making (*35*). Our findings suggest that the same computational principle extends to the coordination of multiple motor effectors. Rather than reflecting noise or interference between vocal and gestural systems, nonlinear mixed selectivity in single neurons could increase the dimensionality of the neural code in AAC and allow downstream circuits to extract commands for multiple effectors from shared inputs (*35*). This framework challenges the idea of specialized vocal circuits (*24*) and provides a possible solution to the problem of flexibly coordinating speech and movement without requiring dedicated, hard-wired pathways for every possible behavioral combination.

In common songbird models such as the zebra finch, the AAC-analog nucleus RA has limited connectivity with surrounding tissue, targeted projections to vocal and respiratory centers in the brainstem and appears to code for vocalizations but not body movement, which is driven by surrounding arcopallial areas not interconnected with RA (*36–39*). Parrot AAC has several anatomical differences from songbird RA, including diffuse boundaries, reciprocal connectivity with surrounding arcopallium and motor thalamus, and diffuse axonal arborizations in the brainstem (*23, 24*). This anatomy more closely resembles the human frontal cortical speech areas, where local circuits may intermix vocal and gestural signals (*25*). We propose that these anatomical differences in AAC, including reciprocal connectivity with local arcopallium, provide a possible substrate for the high-dimensional integration of multiple motor effectors.

The discovery of socially imitated vocal-gestural coupling calls for a re-evaluation of classic vocal learning frameworks in songbirds. The traditional template-comparison model, in which a juvenile matches vocal output to a fixed auditory target through trial and error (*2*, *33*, *40*), does not readily account for the open-ended, socially contingent multimodal imitation observed here. An alternative framework is predictive learning, in which individuals learn to anticipate the multimodal behavior of social partners. Training to forecast upcoming elements in sequential data is sufficient to generate flexible imitative behavior without reliance on fixed templates, as illustrated by contemporary language models (*41*, *42*). Predictive accounts of human communication similarly posit that listeners continuously anticipate what a partner will say and do next (*43*, *44*). Consistent with this idea, behavioral studies show that budgerigars are extremely sensitive to violations of expected syllable sequences during warble song, detecting swapped syllables even after tracking long vocal sequences, suggesting that they actively predict upcoming elements in song (*45*). Because vocal and gestural signals mutually constrain these expectations, predictive frameworks naturally give rise to cross-modal behavioral coupling and mixed neural representations across sensory and motor systems. In humans, the superior temporal sulcus integrates speech sounds, lip movements, facial expressions, and hand gestures, and projects to speech motor areas involved in both comprehension and multimodal production (*43*, *46*). If parrots use human-like predictive learning algorithms to imitate multimodal communication, neural activity in the parrot song system should encode predictions of a social partner’s upcoming vocal or gestural elements during social interactions, and violations of predictions should generate error signals that guide learning. Recordings from AAC and its inputs during listening could test if such predictive multimodal representations exist and, if so, if they are integrated locally within AAC or inherited from distributed forebrain circuits.

Our discoveries support a general neural strategy for complex communication. By integrating vocal and gestural signals within single neurons of a local cortical circuit, the brain may coordinate multiple effectors through a shared code rather than through separate specialized pathways. Such mixing may enable flexibility as new vocalizations and couplings are socially learned and recombined. The budgerigar therefore provides a powerful model for studying the neural basis of learned, embodied communication, a capability that may rely on shared computational principles across avian and human lineages.

## Acknowledgments

We thank members of the Goldberg lab and Eugene Kim and Treasure Nwokoleme for comments and help with data analysis. We thank John Pearson, Miles Martinez, Tim Sainburg, and Richard Hahnloser for discussion on acoustic analysis methods. The authors thank ChatGPT (OpenAI) for assistance with coding suggestions, analysis troubleshooting, and manuscript editing. All scientific decisions, analyses, and interpretations were carried out by the authors.

## Author contributions

Conceptualization: JC, HKT, RC, AR, and JG. designed the study.

Methodology: CJ, HKT, AR, RC, BK, ZZ

Investigation: JC, CJ, AR, RC, BK

Visualization: CJ, HKT

Funding acquisition: JG, IC

Writing – original draft: JG, CJ, HKT

## Data, code, and materials availability

Data and code will be hosted in this <u>OSF.io</u> repository: https://osf.io/2za5b/overview?view_only=e39424109a8247b69374f69019b9b7a9 (a DOI will be generated and added before final publication). Extracted behavioral and neural data (segmentation, labels, spike times, etc.) will be made available in “.m” format (readable by MATLAB and the Python SciPy library), which contains self-describing data structures with human-readable field names. The raw recordings these are based on, while too large to be hosted publicly, can be made available on request. Additional intermediate analysis results will also be made available as .m files. Trained weights for machine learning models used in analysis will be made available as PyTorch .pth files. Python and MATLAB code used to produce the intermediate analysis results and final figures will be made available as well.

## Supplementary Materials

Materials and Methods

Figs. S1 to S6

Table S1

References (1–44)

S1 to S8

## Supplementary Materials for

## Materials and Methods

### Behavioral recordings and social paradigms

For behavioral experiments, two colonies of budgerigars (*Melopsittacus undulatus*) were housed in visual and acoustic isolation (Cohort A, A1-A4; Cohort B, B1-B4) in 18” x 18” x 31.5” cages. Environments were maintained on a 12h light cycle using 850nm IR LEDs to facilitate infrared video acquisition. Mirror-directed behavior was recorded at 90 fps using an IR-sensitive camera (Flir Blackfly S BFS-U3-32S4M, Kowa LM16FC24M lens, Hoya IR (R72) filter) positioned behind a one-way mirror, while synchronized audio was captured at 50 kHz (Audio-Technica A35). For social re-pairing experiments, birds from both cohorts were initially co-housed in a large group cage for 6 weeks before being transitioned into cross-cohort dyads (one bird from each original colony). Dyadic interactions were recorded 10 months post-pairing. Birds were identified by unique facial characteristics (iris contrast, cheek spots)(*47*), and mirror-directed “interactions” were defined as periods of warbling and gesturing exceeding 1 s.

### Warble segmentation

Warble song was segmented and labeled using a fine-tuned WhisperSeg-base transformer model (*14*) trained on approximately 6000 manually labeled syllables from four birds. This model (F1 = 0.878) classified vocalizations into five categories: contact calls, broadband squawks, rhythmic compound syllables, broadband clicks, and G-notes (*10,16*). To quantify acoustic similarity in segmented syllables across individuals, we extracted 11 acoustic features from each syllable spectrogram: the mean and standard deviation of mean frequency, spectral entropy, spectral flatness, spectral skewness, and modulation rate (defined as the first temporal derivative of mean frequency), as well as total syllable duration (*15*). All features were z-score normalized across birds within each syllable label to ensure comparability across feature scales. For each syllable label, we computed pairwise Euclidean distances (d_ij) between the standardized feature vectors of all bird pairs. These distances were transformed into similarity scores using the relation S_ij = 1 / (1 + d_ij). These scores were then averaged for each unique pair of birds. To account for individual differences in self-similarity and vocal variability, factors not typically addressed in standard songbird analyses (*15*), we applied a geometric normalization: each similarity score was divided by the square root of the product of the corresponding self-similarity values, S_ij / sqrt(S_ii * S_jj). We utilized these aggregated, normalized similarity scores to evaluate whether the mean similarity among intra-group pairs differed significantly from inter-group pairs via a permutation test.

## Headbob analyses

In the behavioral dataset, head position (cere) was tracked via DeepLabCut (*22*). Trajectories were detrended using a 340-ms median filter and transformed via a Morlet wavelet (scales 1–60). Headbob bouts were identified by spectral power peaks at scales 40–45, with headbob frequency (F) defined as F = (fc * fs) / a, where fc = pi/3 and fs is the sampling frequency. Temporal coupling between syllables and headbob phase was quantified via bird-level permutation tests (10,000 iterations), using the squared distance between cohort-mean probability curves as a test statistic. Segmented intervals were then manually reviewed to eliminate potential false positives. To quantify syllable production in headbob bias at the level of individual birds, we computed a preference score (Compound − Click) for each bird. Preference scores were positive for all A birds and negative for all B birds, and differed significantly between cohorts (Mann–Whitney U test, N_A=4, N_B=3, p = 0.028), confirming the cohort specific vocal gesture coupling. To quantify coupling of syllable timing to headbob phase, we performed a bird-level permutation test in which phase-dependent probability curves were computed for each animal and birds were randomly reassigned to cohorts (10,000 permutations). The squared distance between cohort-mean curves was used as the test statistic, revealing a significant difference between colonies, p = 0.025). We performed a bird-level permutation test using a global statistic defined as the sum of per-syllable distances between cohort-mean curves. This omnibus test revealed a strong cohort effect (p =9.9×10−5). Post hoc syllable-specific permutation tests with Benjamini–Hochberg FDR correction across five categories showed significant cohort differences for call, click, squawk, and G syllables (q=0.0347 for each), whereas compound syllables approached significance (q=0.0576).

### Latent space analyses and predictive networks

To represent fine-scale subsyllabic acoustic structure, we implemented a shotgun Variational Autoencoder (VAE) with a 32D latent space and tanh activations (*20*). Spectrograms were processed using an 80-ms sliding window (5-ms steps) and resized to 32 x 128 pixels. The model was trained for 50 epochs (ADAM optimizer, learning rate 0.001) using a 90/10 train-test split. Cross-bird acoustic similarity was quantified using a reconstruction-based metric. For a VAE trained on a reference bird, mean-squared reconstruction error (MSE) was computed for both training data and evaluation data from held out data or another bird. Similarity was defined as the ratio MSE(train)/MSE(eval), with values approaching 1 indicating high similarity. For supplemental analyses, similarity was also assessed using conventional acoustic features (frequency statistics, entropy, spectral shape, modulation rate, and duration)(*15*). Feature vectors were z-scored within syllable category, pairwise Euclidean distances were computed, and distances were converted to similarity scores. Group differences were evaluated by permutation testing. An LSTM classifier was trained to predict head-bob occurrence based on the preceding 200 ms of VAE latent features. The network consisted of two LSTM layers (hidden size 50) followed by a fully connected output layer. Models were trained per bird (80% training / 20% testing) using cross-entropy loss and evaluated on held-out and cross-bird data. The output log-likelihood was smoothed using a 100 millisecond moving average. The learning rate was 0.01, batch size 20, and the model was trained for 50 epochs. Classifier performance was evaluated on both held-out and cross-bird datasets.

### Surgery and neural recording

For chronic neural recordings, 4 birds were anesthetized with isoflurane inhalation and 16-channel moveable wire bundles were implanted into AAC coordinates (+5.85-6.05 A, +5.4 L [relative to lambda], 2.6-3.0 V [relative to pial surface]; head angle 30°). Birds were then placed in dyadic chambers with a partner for post-operative recovery on a 12 hour light/dark cycle with ad libitum food and water. Neural signals were amplified, filtered, and digitized on Intan RHD headstages equipped with 3-axis accelerometers. Headstages were connected to the implanted 16-channel moveable electrode bundles from Innovative Neurophysiology. Electrodes were extended or retracted on a semi-daily basis by manual turning of a screw in the morning before the enclosure lights came on. Neural signals were further band-pass filtered and sorted in a custom MATLAB GUI.

### Neural spiking alignment to vocalizations and headbobs

To test if a neuron exhibited a significant response to a given syllable type, spike times were aligned to syllable onset and binned in 10 ms intervals from −0.1 s relative to onset through the median duration of that syllable type. Peri-stimulus time histograms (PSTHs) were computed by averaging across trials and smoothed using a 3-bin moving average. Headbob-aligned PSTHs were constructed using the same procedure. To assess statistical significance of observed firing rate modulations, we generated a null distribution using circular spike-time shuffling (*48*). For each neuron and syllable type, spike trains were randomly circularly shifted 1000 times, and a shuffled PSTH was computed for each iteration. The maximum and minimum firing rates were extracted from each shuffled PSTH to form null distributions of peak and trough activity. The empirical maximum and minimum firing rates were then compared to these null distributions to compute p-values. A neuron was considered to exhibit a significant vocal response if either the maximum or minimum firing rate exceeded the 95th percentile of the corresponding null distribution (p < 0.05). Headbob-aligned PSTHs were constructed using the same procedure. Neurons recorded during at least 20 headbobs with at least 20 spikes during those headbobs were included in the headbob-related analyses in Fig 3.

### Headbob influence on syllable-aligned neural activity

For neurons that exhibited significant call-aligned activity, all call syllables were time-warped to the median duration and grouped according to whether they occurred during headbobs (HB) or outside of them. To isolate neural modulation attributable to the HB context, each HB syllable was paired with its nearest acoustic neighbor produced outside of HB (cosine distance < 0.3 in flattened spectrogram space). Only HB syllables with a matched non-HB counterpart were included in the analysis. Time-warped spike trains from HB syllables and their acoustically matched non-HB counterparts were used to construct firing rate histograms for the two conditions. To assess statistical significance of firing rate differences in each 10 ms time bin, syllables and associated spike trains were randomly reassigned pseudo-labels (HB or non-HB), and firing rate histograms were recomputed for 10,000 iterations to generate a null distribution of bin-wise differences. Observed differences were compared against this null distribution. Neurons exhibiting a significant difference (p<0.01) in at least 3 consecutive time bins were considered to be HB-modulated.

### Statistical analysis of effect of recording depth on neural representations

To assess population-level effects of recording depth on neural activity while accounting for hierarchical data structure, we combined principal component analysis (PCA) with linear mixed-effects modeling (LME) and a within-channel permutation test. Firing rate matrices were first reduced using PCA (units x time bins). PCs explaining at least 95% of the cumulative variance were retained for all subsequent analysis. Each retained PC was weighted by its proportion of explained variance to preserve the contribution of dominant population modes. For each retained PC, we fit a linear mixed-effects model with both linear and quadratic depth terms:

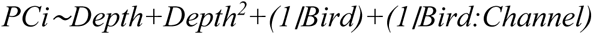

Depth was mean-centered within-recording channel prior to analysis to isolate within-channel depth structure and eliminate between-channel offsets. Random intercepts for Bird and Bird x Channel accounted for repeated measurements and nested recording structure. Linear and quadratic fixed-effect coefficients were extracted for each PC. A single population-level depth effect magnitude was computed as the variance-weighted euclidean norm across PCs:

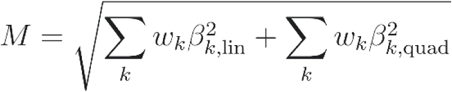

Where w_k_ is the variance explained by PC *k*. Statistical significance was assessed using a nonparametric permutation test. Depth values were randomly permuted within-channel, preserving channel-specific depth distributions while breaking the mapping between depth and neural activity. For each permutation, the full PCA–LME procedure was repeated and a null distribution of population magnitudes was constructed. The p-value was computed as the proportion of null magnitudes greater than or equal to the observed value.

### Pitch tuning and acoustic feature GLM

For each neuron, for all segmented syllables, a ‘harmonic index’ was calculated every 2 ms by multiplying the harmonic ratio (MATLAB 2022a Audio Toolbox, Mathworks) by a Fundamental Index (FI), defined as the prominence of the fundamental frequency estimated from autocorrelation-based pitch tracking. Pitch, entropy, amplitude, and frequency modulation was computed using Sound Analysis Pro (SAP) (15) in 2 ms bins, and neural spikes were counted in the time window −30 to −5ms for each bin. Every bin with HR x FI > 0.6 was included in the pitch analysis. Relative contributions of the acoustic features to neuronal spiking was assessed using a Poisson generalized linear model (GLM) with an exponential link function. GLM significance (p<0.01) for each model was calculated using the likelihood ratio test statistic and shuffled permutations as described. Relative contribution of each predictor was calculated using McFadden’s R2:

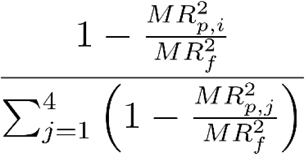

Where MR^2^_p,i_ is McFadden’s R2 for the partial model with the i’th predictor excluded and MR^2^_f_ corresponds to the full model. The above analyses precisely replicate procedures described in a previous paper linking AAC spiking to acoustic features (30).

Nonlinear encoding models of vocal and movement contributions to neural activity

For each neuron with at least 20 headbob renditions, we modeled mean firing rate in sliding 50 ms windows as a function of vocal features, movement features, or both.

Spectrograms spanning each headbob were processed using a 50 ms window (3 ms step) and passed through a trained autoencoder to obtain a 32-dimensional latent representation of vocal acoustics (*20*). In parallel, movement features were extracted within the same windows from skull-mounted accelerometers. Velocities along dorsal–ventral and anterior–posterior axes were estimated by cumulative trapezoidal integration followed by median filtering. For each window, velocity values at the beginning and end of the window were computed along both axes.

Headbob phase at the beginning and end of the window, as well as absolute and relative time within the headbob bout, were included, yielding eight movement features per window. To capture nonlinear interactions between vocal and kinematic variables, we used gradient-boosted regression trees (XGBoost) (*26*, *27*). For each neuron, an ensemble of 200 trees (maximum depth 10) was trained with L1 (α = 0.1) and L2 (λ = 5) regularization. Models were evaluated using grouped k-fold cross-validation, in which 20 s recording files served as independent groups, ensuring that all time bins from a given file were held out together to prevent inflation of performance estimates due to temporal autocorrelation (*49,50*). Model performance was quantified as reduction in mean squared error relative to a null model predicting the mean firing rate of the training data. For each held-out file, improvement relative to the null was computed. Statistical significance was assessed using a one-sided Wilcoxon signed-rank test across files. To dissociate vocal and movement contributions, partial models using only vocal or only movement predictors were compared to the full model. Improvement of the full model over each partial model was evaluated using the same nonparametric test across held-out files. Neurons were classified as mixed if the full model significantly outperformed both partial models, indicating that joint inclusion of acoustic and kinematic features was required to explain spiking activity.

## Movies S1 – S8

Example video (0.5× speed) of a budgerigar performing headbobs toward a one way mirror. Shown alongside the video are the vocal spectrogram (top), head velocity (middle), and projections of the latent acoustic space with relative time in the headbob sequence and cycle phase overlaid (bottom right).

---

**Fig. S1.**
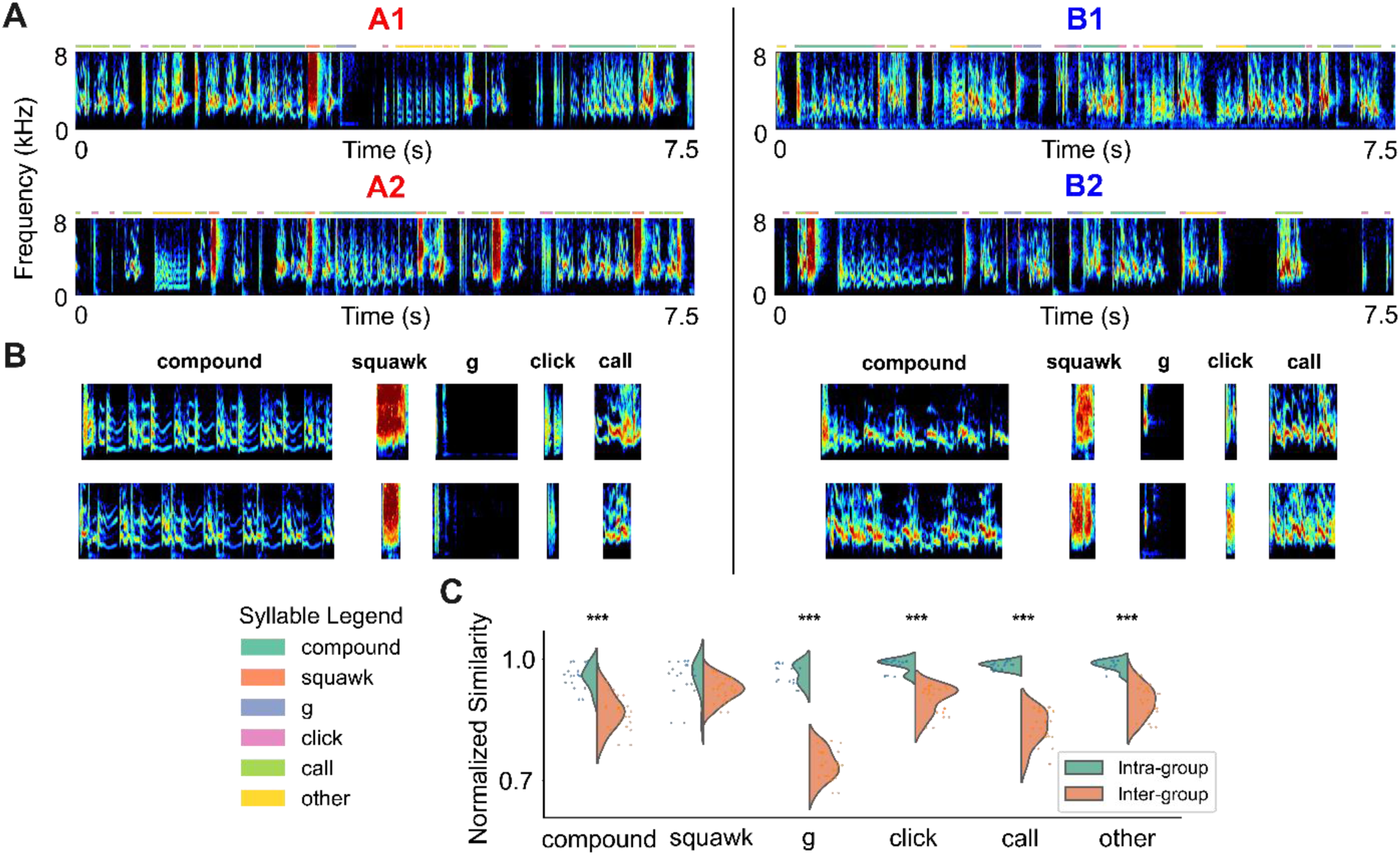
Conventional syllable segmentation and similarity scoring methods confirm imitation by cagemates. (**A**) Example warble bouts from representative birds in Cohort A (A1, A2) and Cohort B (B1, B2), color-coded by syllable category. (**B**) Representative spectrograms for each syllable type: Compound (long duration syllables with complex intrasyllabic repetitive acoustic structure); Squawk (brief broadband note); ‘G’ (click followed seamlessly by low frequency tail); Click (broadband note <50 ms); Call (contact call inserted into warble song). ‘Other’ was a category of other highly variable syllables unrecognizable to human eye or WhisperSeg. (**C**) Normalized similarity distributions comparing intra-versus inter-group pairs for each syllable class (***p< 0.001, two-sample t-test), indicating stronger convergence within cohorts.

**Fig. S2.**
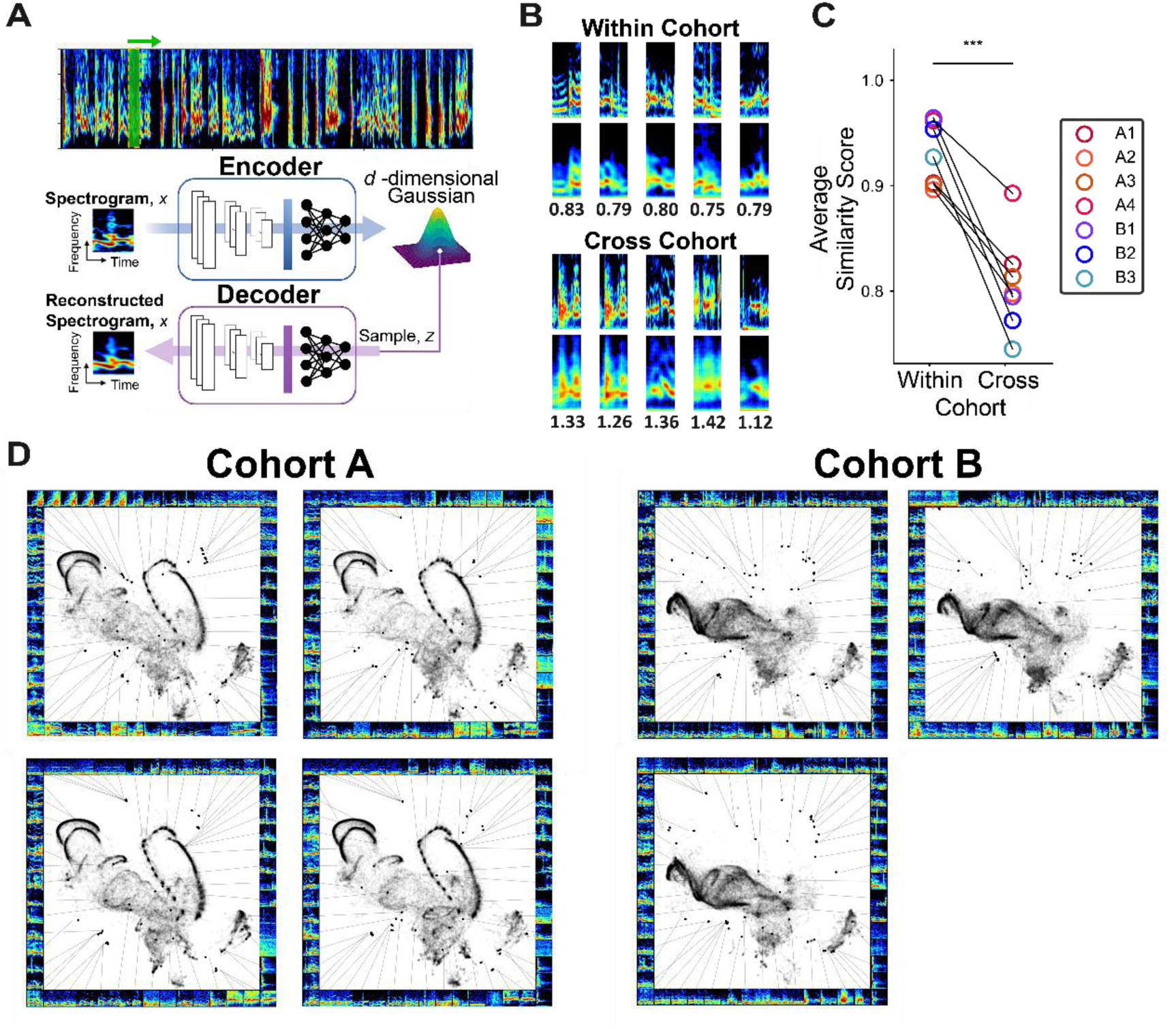
**VAE latent space analyses quantify warble imitation**. (**A**) A “shotgun” variational autoencoder (VAE) approach in which sliding 80-ms spectrogram windows (5-ms step size) are embedded into a 32-dimensional latent space. (**B**) Example reconstructions and reconstruction error for a VAE trained on bird A1, evaluated on held-out A1 warbles (top) or warbles from bird B1 (bottom). Reconstruction error is lower for within-bird evaluation and higher for cross-cohort evaluation, reflecting acoustic dissimilarity. (**C**) Acoustic similarity was quantified using a reconstruction-based similarity score, defined as the ratio of mean-squared reconstruction error (MSE) on the training data to the MSE on an evaluation dataset. Similarity scores approach 1 for acoustically similar warbles and decrease as vocal structure diverges; scores were significantly higher within cohorts than across cohorts. (**D**) Two-dimensional UMAP projections of the VAE latent space for birds A1–A4 and B1–B3. Warble manifolds are highly similar within cohorts but distinct across cohorts, revealing socially learned, colony-specific acoustic structure.

**Fig. S3.**
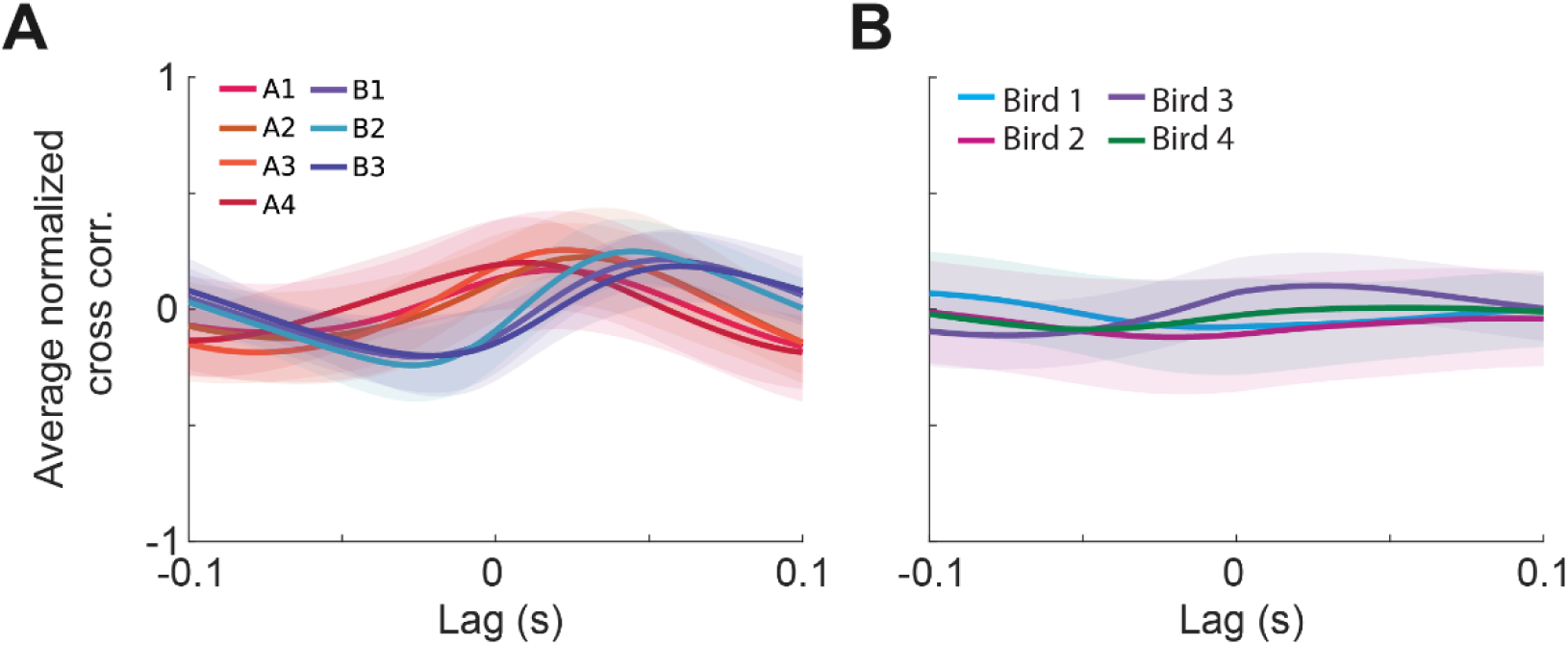
**The relationship between head velocity and sound amplitude was colony-specific**. **(A)** Average +/1 STD cross correlation between sound amplitude and head velocity. A permutation test showed that cohort A and B birds exhibited significantly distinct curve shapes (p<0.01). **(B)** Data plotted as in A for the 4 birds examined in the electrophysiology experiments, which had no social contact with the colony A or B birds.

**Fig. S4.**
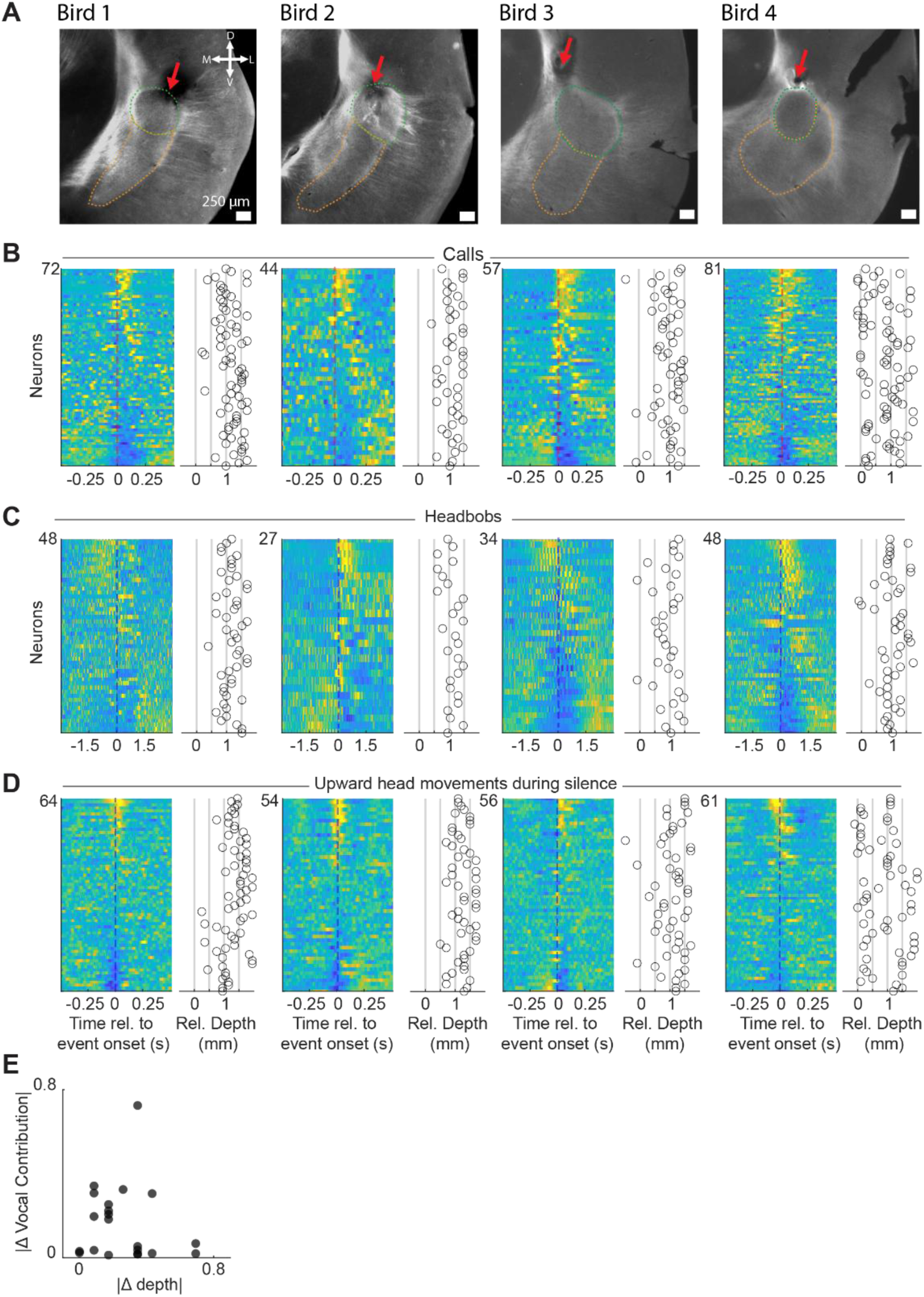
Vocal and gestural representations are spatially overlapping in AAC. (**A**) Histological confirmations of cannula position (red arrows), dorsal to AAC core, from which electrodes were advanced. Green and orange dotted lines denote boundaries of dorsal core and ventral shell, respectively. (**B**) Call-aligned z-scored firing rates of 72 AAC neurons sorted by the first principal component (PC1) of the individual bird population call-aligned activity; scatter plots show the recording depth of each neuron (row-aligned), relative to the estimated start of AAc core (mm). (**C**) Data plotted as in B for headbob aligned activity. In all birds, call- and headbob-locked neural representations were found spatially intermingled at all sites. (**D**) Data plotted as in B,C for activity aligned to upward movements of the head during silent times. (**E**) Within bird, within channel comparisons of absolute difference in recording depth vs absolute difference in the vocal contribution metric shown in Fig. 4G.

**Fig. S5.**
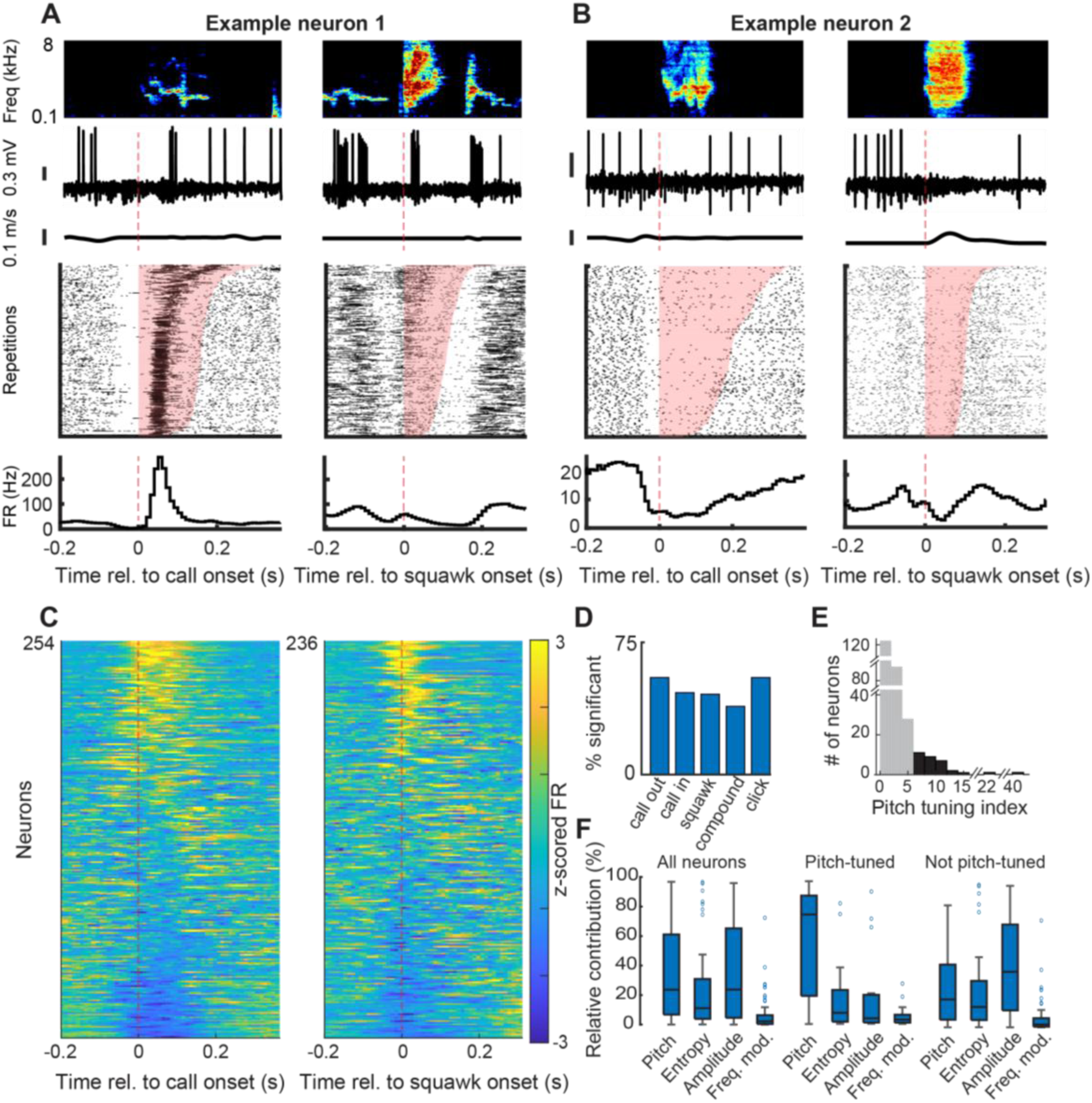
**Acoustic and segmentation-based analyses confirm vocalization-locked activity of AAC neurons**. (**A-B**) Call (left) and squawk (right) aligned activity of two example AAC neurons. (**C**) Population call- and squawk-aligned activity for 254 and 236 AAC neurons, respectively, sorted by the first principal component (PC1) score of the call- or squawk aligned population response. (**D**) Proportion of neurons exhibiting significant firing-rate maxima or minima for different vocalization types. (**E**) Distribution of pitch-tuning index values. Gray bars denote non-pitch-tuned neurons (index < 6); black bars denote pitch-tuned neurons (index ≥ 6). Axes are broken to enhance visualization. (**F**) Relative contributions of different acoustic features to firing-rate prediction in a generalized linear model (GLM). These findings confirm a recent study by the Long lab (*25*), including neuronal tuning to pitch, showing that the gestural signals we observe are in the same population of neurons.

**Fig. S6.**
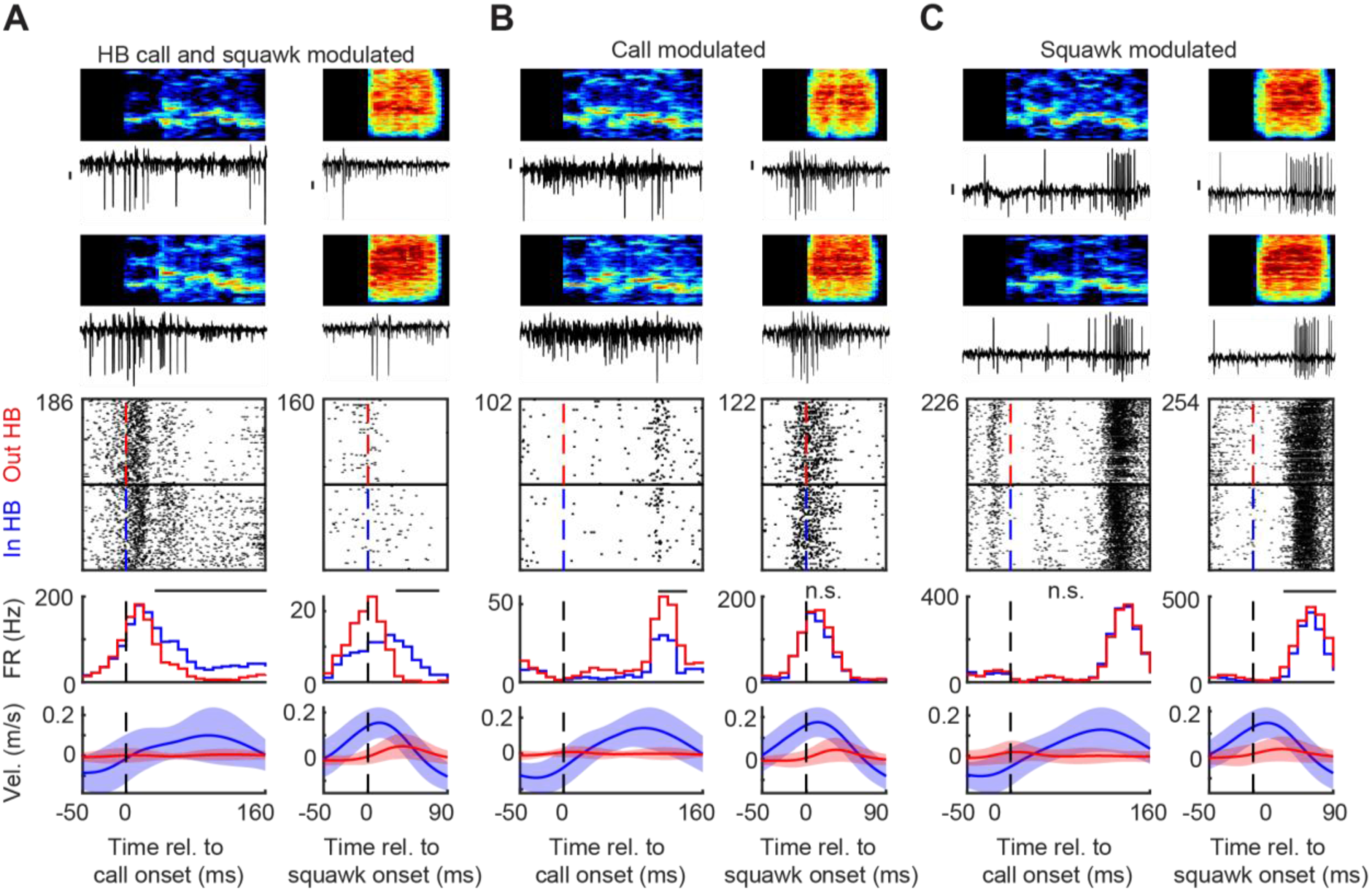
**Headbobs can modulate syllable-locked AAC activity even when syllable acoustics are matched**. (**A**) Neuron showing differential activity for calls produced outside versus during headbob (HB) bouts. Left column (top to bottom): spectrograms and spiking for nearest-neighbor call renditions outside and inside HB; call-aligned rasters (outside HB, top; inside HB, bottom); rate histogram for outside (red) and inside (blue) HB trials, and call-aligned head velocity traces (mean±SD.). Right column: same analyses for squawk syllables. Neurons were classified as HB-modulated if ≥3 consecutive 10-ms bins differed significantly between conditions (nonparametric shuffle test, p < 0.001, black bars). (**B**) Neuron exhibiting significant HB-dependent modulation for calls only. (**C**) Neuron exhibiting HB-dependent modulation for squawks only.

**Table S1.**
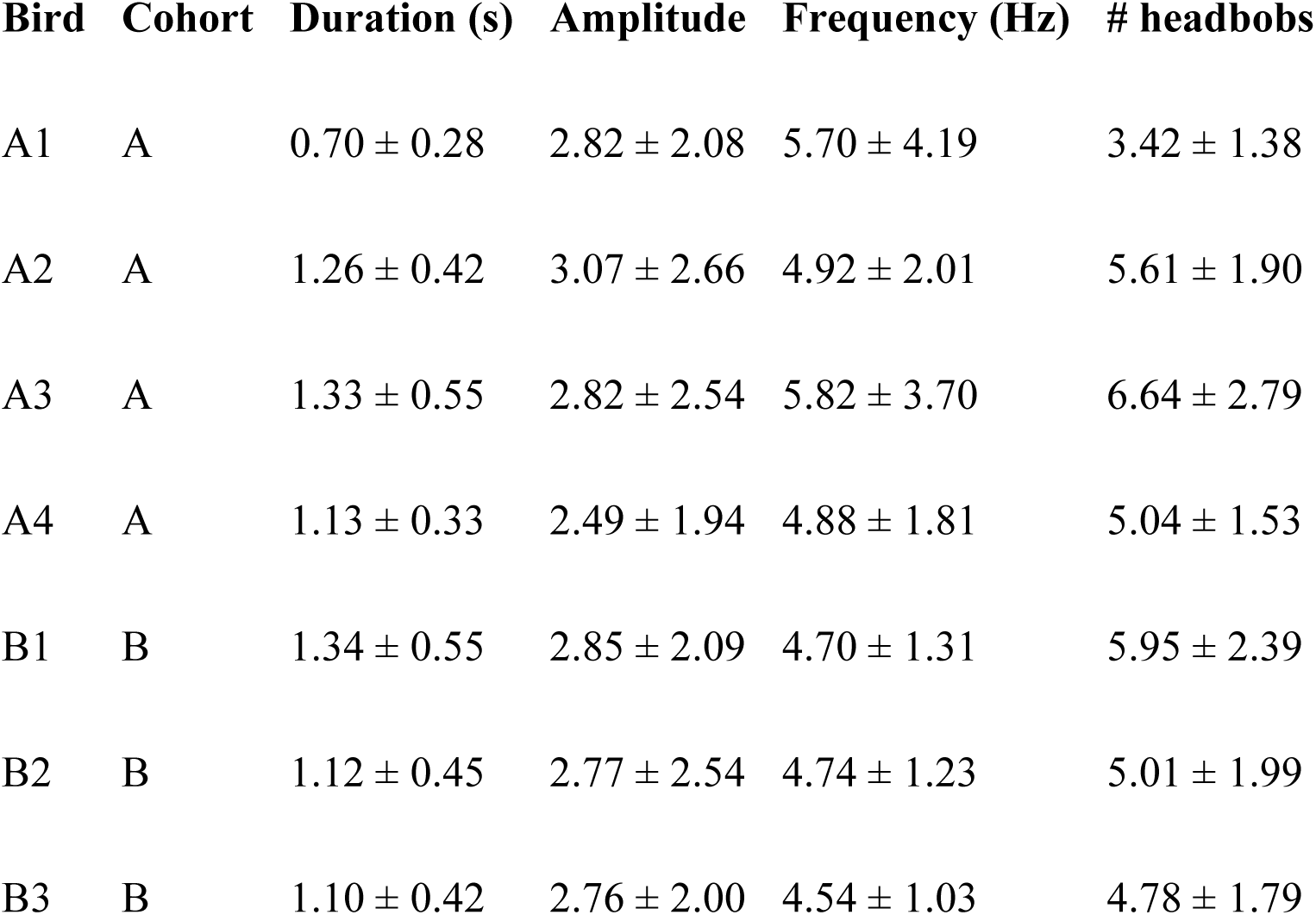
Headbob kinematic measures by bird. Values represent per-bird medians across headbob bouts for bout duration, amplitude, bob frequency, and number of bobs per bout. Birds are grouped by cohort (A or B).

## References

1. S. Kita, Cross-cultural variation of speech-accompanying gesture: A review. Lang. Cogn. Process. 24, 145–167 (2009).

2. E. L. Mackevicius, M. S. Fee, Building a state space for song learning. Curr. Opin. Neurobiol. 49, 59–68 (2018).

3. R. Mooney, Birdsong. Curr. Biol. 32, R1090–R1094 (2022).

4. S. R. Partan, P. Marler, Issues in the classification of multimodal communication signals. Am. Nat. 166, 231–245 (2005).

5. Ç. Akçay, M. D. Beecher, Multi-modal communication: song sparrows increase signal redundancy in noise. Biol. Lett. 15, 20190513 (2019).

6. N. Ota, M. Gahr, M. Soma, Tap dancing birds: the multimodal mutual courtship display of males and females in a socially monogamous songbird. Sci. Rep. 5, 16614 (2015).

7. J. P. Higham, Multimodal communication in nonhuman primates and humans. Annu. Rev. Anthropol. 54, 427–441 (2025).

8. L. Fusani, R. E. Hutchison, J. B. Hutchison, Vocal-postural co-ordination of a sexually dimorphic display in a monomorphic species: The Barbary dove. Behaviour 134, 321–335 (1997).

9. B. G. Cooper, F. Goller, Multimodal signals: enhancement and constraint of song motor patterns by visual display. Science 303, 544–546 (2004).

10. N. Wewhare, A. Krishnan, Individual-specific associations between warble song notes and body movements in budgerigar courtship displays. Biol. Open 13 (2024).

11. B. F. Brockway, Ethological studies of the budgerigar: reproductive behavior. Behaviour 23, 294–323 (1964).

12. A. D. Patel, J. R. Iversen, M. R. Bregman, I. Schulz, Studying synchronization to a musical beat in nonhuman animals. Ann. N. Y. Acad. Sci. 1169, 459–469 (2009).

13. S. M. Farabaugh, E. D. Brown, R. J. Dooling, Analysis of warble song of the budgerigarmelopsittacus undulatus. Bioacoustics 4, 111–130 (1992).

14. N. Gu, K. Lee, M. Basha, S. K. Ram, G. You, R. Hahnloser, Positive transfer of the whisper speech transformer to human and animal Voice Activity Detection, bioRxiv (2023). 10.1101/2023.09.30.560270.

15. O. Tchernichovski, F. Nottebohm, C. E. Ho, B. Pesaran, P. P. Mitra, A procedure for an automated measurement of song similarity. Anim. Behav. 59, 1167–1176 (2000).

16. H.-W. Tu, E. W. Smith, R. J. Dooling, Acoustic and perceptual categories of vocal elements in the warble song of budgerigars (Melopsittacus undulatus). J. Comp. Psychol. 125, 420–430 (2011).

17. A. J. Madabhushi, N. Wewhare, P. Binwal, V. Agarwal, A. Krishnan, Higher-order dialectic variation and syntactic convergence in the complex warble song of budgerigars. J. Exp. Biol. 226, jeb245678 (2023).

18. D. C. Mann, W. Fitch, H.-W. Tu, M. Hoeschele, Universal principles underlying segmental structures in parrot song and human speech. Sci. Rep. 11 (2021).

19. T. Sainburg, M. Thielk, T. Q. Gentner, Finding, visualizing, and quantifying latent structure across diverse animal vocal repertoires. PLoS Comput. Biol. 16, e1008228 (2020).

20. J. Goffinet, S. Brudner, R. Mooney, J. Pearson, Low-dimensional learned feature spaces quantify individual and group differences in vocal repertoires. Elife 10 (2021).

21. L. McInnes, J. Healy, J. Melville, UMAP: Uniform Manifold Approximation and Projection for Dimension Reduction, arXiv [stat.ML] (2018). http://arxiv.org/abs/1802.03426.

22. A. Mathis, P. Mamidanna, K. M. Cury, T. Abe, V. N. Murthy, M. W. Mathis, M. Bethge, DeepLabCut: markerless pose estimation of user-defined body parts with deep learning. Nat. Neurosci. 21, 1281–1289 (2018).

23. G. Feenders, M. Liedvogel, M. Rivas, M. Zapka, H. Horita, E. Hara, K. Wada, H. Mouritsen, E. D. Jarvis, Molecular mapping of movement-associated areas in the avian brain: a motor theory for vocal learning origin. PLoS One 3, e1768 (2008).

24. M. Chakraborty, E. D. Jarvis, Brain evolution by brain pathway duplication. Philos. Trans. R. Soc. Lond. B Biol. Sci. 370 (2015).

25. R. L. Tatarsky, Z. Guo, S. C. Campbell, H. Kim, W. Fang, J. T. Perelmuter, E. R. Schuppe, K. W. Conway, H. K. Reeve, A. H. Bass, Acoustic and postural displays in a miniature and transparent teleost fish, Danionella dracula. J. Exp. Biol. 225 (2022).

26. J. M. Wild, J. F. Botelho, Involvement of the avian song system in reproductive behaviour. Biol. Lett. 11, 20150773 (2015).

27. W. C. Lancaster, H. J. O. W., A. W. Keating, Respiratory muscle activity in relation to vocalization in flying bats. Journal of Experimental Biology 198, 175–191 (1995).

28. G. F. Striedter, The vocal control pathways in budgerigars differ from those in songbirds. J. Comp. Neurol. 343, 35–56 (1994).

29. M. Chakraborty, S. Walloe, S. Nedergaard, E. E. Fridel, T. Dabelsteen, B. Pakkenberg, M. F. Bertelsen, G. M. Dorrestein, S. E. Brauth, S. E. Durand, E. D. Jarvis, Core and Shell Song Systems Unique to the Parrot Brain. PLoS One 10, e0118496 (2015).

30. Z. Yang, M. A. Long, Convergent vocal representations in parrot and human forebrain motor networks. Nature 640, 427–434 (2025).

31. T. Chen, C. Guestrin, “XGBoost: A Scalable Tree Boosting System” in Proceedings of the 22nd ACM SIGKDD International Conference on Knowledge Discovery and Data Mining (ACM, New York, NY, USA, 2016; 10.1145/2939672.2939785).

32. A. S. Benjamin, H. L. Fernandes, T. Tomlinson, P. Ramkumar, C. VerSteeg, R. H. Chowdhury, L. E. Miller, K. P. Kording, Modern machine learning as a benchmark for fitting neural responses. Front. Comput. Neurosci. 12, 56 (2018).

33. R. Chen, J. H. Goldberg, Actor-critic reinforcement learning in the songbird. Curr. Opin. Neurobiol. 65, 1–9 (2020).

34. J. A. Tourville, F. H. Guenther, The DIVA model: A neural theory of speech acquisition and production. Lang. Cogn. Process. 26, 952–981 (2011).

35. S. Fusi, E. K. Miller, M. Rigotti, Why neurons mix: high dimensionality for higher cognition. Curr. Opin. Neurobiol. 37, 66–74 (2016).

36. J. M. Wild, Descending projections of the songbird nucleus robustus archistriatalis. J. Comp. Neurol. 338, 225–241 (1993).

37. R. C. Yuan, S. W. Bottjer, Multidimensional tuning in motor cortical neurons during active behavior. eNeuro 7, ENEURO.0109–20.2020 (2020).

38. S. J. Sober, M. J. Wohlgemuth, M. S. Brainard, Central contributions to acoustic variation in birdsong. J. Neurosci. 28, 10370–10379 (2008).

39. A. Leonardo, M. S. Fee, Ensemble coding of vocal control in birdsong. J. Neurosci. 25, 652–661 (2005).

40. P. Marler, A comparative approach to vocal learning: Song development in white-crowned sparrows. J. Comp. Physiol. Psychol. 71, 1–25 (1970).

41. E. M. Bender, T. Gebru, A. McMillan-Major, S. Shmitchell, “On the dangers of stochastic parrots: Can language models be too big?” in Proceedings of the 2021 ACM Conference on Fairness, Accountability, and Transparency (ACM, New York, NY, USA, 2021; 10.1145/3442188.3445922).

43. A. Vaswani, N. Shazeer, N. Parmar, J. Uszkoreit, L. Jones, A. N. Gomez, L. Kaiser, I. Polosukhin, Attention is all you need, arXiv [cs.CL] (2017). http://arxiv.org/abs/1706.03762.

43. J. I. Skipper, S. Goldin-Meadow, H. C. Nusbaum, S. L. Small, Speech-associated gestures, Broca’s area, and the human mirror system. Brain Lang. 101, 260–277 (2007).

44. S. Garrod, M. J. Pickering, “Interactive alignment and prediction in dialogue” in Advances in Interaction Studies (John Benjamins Publishing Company, Amsterdam, 2013), pp. 193–204.

45. H.-W. Tu, A. R. Fishbein, W. J. Idsardi, R. J. Dooling, Perception of sequential structure in budgerigar (Melopsittacus undulatus) warble song. J. Acoust. Soc. Am. 158, 2863–2868 (2025).

46. J. H. Venezia, K. I. Vaden Jr, F. Rong, D. Maddox, K. Saberi, G. Hickok, Auditory, visual and audiovisual speech processing streams in superior temporal sulcus. Front. Hum. Neurosci. 11, 174 (2017).

47. S. D. Brown, R. J. Dooling, Perception of conspecific faces by budgerigars (Melopsittacus undulatus): II. Synthetic models. J. Comp. Psychol. 107, 48–60 (1993).

48. J. H. Goldberg, A. Adler, H. Bergman, M. S. Fee, Singing-related neural activity distinguishes two putative pallidal cell types in the songbird basal ganglia: comparison to the primate internal and external pallidal segments. J. Neurosci. 30, 7088–7098 (2010).

49. K. Hardcastle, N. Maheswaranathan, S. Ganguli, L. M. Giocomo, A multiplexed, heterogeneous, and adaptive code for navigation in medial entorhinal cortex. Neuron 94, 375–387.e7 (2017).

50. W. Truccolo, U. T. Eden, M. R. Fellows, J. P. Donoghue, E. N. Brown, A point process framework for relating neural spiking activity to spiking history, neural ensemble, and extrinsic covariate effects. J. Neurophysiol. 93, 1074–1089 (2005).

